# Four decades of phenology in an alpine amphibian: trends, stasis, and climatic drivers

**DOI:** 10.1101/2022.08.16.503739

**Authors:** Omar Lenzi, Kurt Grossenbacher, Silvia Zumbach, Beatrice Lüscher, Sarah Althaus, Daniela Schmocker, Helmut Recher, Marco Thoma, Arpat Ozgul, Benedikt R. Schmidt

## Abstract

1. Strong phenological shifts in response to changes in climatic conditions have been reported for many species, including amphibians, which are expected to breed earlier. Phenological shifts in breeding are observed in a wide number of amphibian populations, but less is known about populations living at high elevations, which are predicted to be more sensitive to climate change than lowland populations.
2. The goal of this study is to assess the main factors determining the timing of breeding in an alpine population of the common toad (*Bufo bufo*) and to describe the observed shifts in its breeding phenology.
3. We modelled the effect of environmental variables on the start and peak dates of the breeding season using 39 years of individual-based data. In addition, we investigated the effect of the lunar cycle, as well as the individual variation in breeding phenology. Finally, to assess the individual heterogeneity in the timing of breeding, we calculated the repeatability of the timing of arrival at the breeding site.
4. Breeding advanced to earlier dates in the first years of the study but the trend continued only until the mid 1990s, and stabilised afterwards. Overall, toads are now breeding on average around 30 days earlier than at the start of the study period. High temperatures and low snow cover in winter and spring, as well as reduced spring precipitation were all associated with earlier breeding. Additionally, we found evidence of males arriving on average before females at the breeding site but no clear and strong effect of the lunar cycle. We only found weak evidence of among-individual variation in shifts in the breeding phenology, as well as a low repeatability of arrival timing.
5. Our findings show that the observed changes in breeding phenology are strongly associated with the environmental conditions. These results contribute to filling a knowledge gap on the effects of climate change on alpine amphibian populations. Moreover, we show that changes in phenology, especially in the mountains, can be hard to predict as local microclimatic conditions do not necessarily reflect the observed global climatic trends.

## Introduction

Phenology refers to the timing of periodical events (e.g., seasonal migration, spring flowering) in relation to biotic and abiotic factors, and is a key element of the life cycle in a multitude of organisms. Phenology is normally determined by a combination of a genetic and an environmental component (Quinn & Wetherington, 2002; Tang et al., 2016). Thus, climate change can shift the phenology of many species, potentially leading to mismatches between demand and availability of resources (Parmesan & Yohe, 2003; Visser & Gienapp, 2019; Iler et al., 2021). These shifts can have large effects on the demography of populations, as individuals cannot benefit from the optimal conditions at the right time, with consequences on their fitness (Visser & Gienapp, 2019; Iler et al., 2021). Therefore, it is important to describe and quantify phenological shifts and their causes.

Phenology has a key role in amphibians as well, especially in species living in temperate regions, where various aspects of the annual cycle are determined by seasonality (Duellman & Trueb, 1986; Gotthard, 2001; Hartel et al., 2007). The environmental component is more important than the genetic component in explosive breeders (*sensu* Wells, 1977). In fact, explosive breeders reproduce once a year around springtime and the timing is linked to specific environmental signals such as increasing day length, temperature, and rainfall, which can trigger the migration of amphibians from the hibernation sites to the breeding ponds (Semlitsch, 1985; Oseen & Wassersug, 2002; While & Uller, 2014; Ficetola & Maiorano, 2016). Other important environmental factors affecting the timing of breeding in explosive breeders can be the lunar cycle (Grant et al., 2009; Green et al., 2016; Arnfield et al., 2012; Jarvis et al., 2021) or the hydrological cycle of breeding ponds (Semlitsch et al., 1993). Previous studies have also identified a possible genetic component in triggering the migration to the breeding site and thus the start of the breeding season (Heusser & Ott, 1968; Semlitsch et al., 1993; Phillimore et al., 2010). Breeding phenology also shows individual variation, as the animals will arrive at different times at the breeding site. The causes of individual-level variation are multifold and can include genetics (Heusser & Ott, 1968; Semlitsch et al., 1993), sex and size (Loman & Madsen, 1986), body condition (Kokko, 1999) as well as features of the hibernation site, such as distance from the breeding site, which in *Bufo bufo* can be up to more than 1000 m (Sztatecsny & Schabetsberger, 2005; Kovar et al., 2009).

While most studies on amphibians across species and locations have found earlier spring breeding in response to climate change (Beebee, 1995; Blaustein et al., 2001; Parmesan, 2007; While & Uller, 2014), phenological delays have also been observed (e.g., Arnfield et al., 2012; Arietta et al., 2020). In other cases, non-linear responses to environmental drivers such as the North Atlantic Oscillation were observed (Prodon et al., 2020). The direction and magnitude of phenological shifts are therefore variable among and within species, as they can depend on the specific environmental conditions that the populations are experiencing at the local scale, or on the genetic structure of said populations (Phillimore et al., 2010; Bison et al., 2021).

Shifts in phenology can have adverse effects on amphibians, as phenological mismatches can affect predator-prey dynamics and food availability (Todd et al.,2011; Reinhardt et al., 2015; Jara et al., 2019; Visser & Gienapp, 2019). In temperate regions, early breeding can expose eggs and hatched tadpoles more frequently to late frost events, thus increasing mortality (Muir et al., 2014; Bison et al., 2021). On the other hand, in the absence of frost or drying events, earlier breeding might be beneficial as it allows post-metamorphic toadlets more time to fully develop in summer before hibernation (Reading & Clarke, 1999; Reading, 2010). Delayed breeding can also have a negative outcome on the population, as it can result in increased mortality in juveniles that could not fully grow before their first hibernation (Morin et al., 1990; Garner et al., 2011; Sinsch & Schäfer, 2016). Even though this phenomenon can be compensated in some cases with an accelerated growth rate, this can come at the cost of reduced defences against predation (Orizaola et al., 2016). Thus, phenological shifts and their causes should be identified and better understood, as they can help design and prioritise conservation and management actions.

The consequences of phenological shifts could be exacerbated in ecosystems less resilient to climate change. Mountains are among the most threatened ecosystems (Thompson, 2000; Diaz et al., 2003, but see Körner & Hiltbrunner, 2021) and are predicted to warm more rapidly in the northern hemisphere (Nogués-Bravo et al., 2007; Keiler et al., 2010; Vitasse et al., 2021). The phenology of plant and animal populations at high elevations is shifting on average towards earlier dates (Vitasse et al., 2021). Long-term studies on amphibian populations living at high elevations are scarce, and not much is known about how their breeding phenology is changing. These populations experience different environmental conditions (e.g., increased amount of snow and colder temperatures) compared to their lowland counterparts. Thus, different environmental variables potentially play a bigger role in determining breeding phenology compared to what is observed at lower elevations (Nufio et al., 2010; Bison et al., 2020).

Using 39 years of data on an explosive-breeding amphibian population living at a high elevation (*B. bufo*), we study the relationship between breeding phenology and the environment. More specifically, our goal is to (i) identify the environmental variables (e.g., temperature, snow cover, moon cycle) that could be driving the observed breeding phenology of this population (both the start and the peak of the breeding season), (ii) analyse if there is significant variation in the phenological shifts among individuals, (iii) obtain a measure of individual heterogeneity, by calculating individual-level repeatability (i.e., upper limit of heritability; Falconer, 1981; Lessells & Boag, 1987; Semlitsch et al., 1993) of the timing of arrival at the breeding site for both males and females.

## Material and Methods

### Life-history data

The study site is a pond located above Grindelwald, below the Grosse Scheidegg mountain pass (canton of Bern, Switzerland, 46.65240 N, 8.09683 E), at an elevation of 1841 m a.s.l. The pond measures approximately 10 m x 30 m, with a maximum depth of about 1 m. Since 1982, we have captured annually all the toads that come to breed at the study pond. We then marked (first by toe-clipping, then starting in 1993 by implanting PIT tags), measured, and released them in the same place (Hemelaar, 1988; Grossenbacher, 2002). To make sure we captured both early and late arrivers, we repeated this procedure for on average 5–6 nights, with breaks in-between of about 2-4 days (i.e, the data conform to Pollock’s (1982) robust design). The length of the fieldwork period usually covers the breeding season duration, which typically lasts about two weeks at our study pond. This design also had the advantage of not overly stressing the toads. In total, for the period 1982–2020, 3053 uniquely recognizable individuals have been caught, of which 1852 were males and 1201 females. For each individual we have a record of presence for each capture night over the study period. Given the reduced size of the pond and the repeated capture rounds within a capture night, we assumed high capture probabilities (capture probability p ≈ 0.85 per year based on a preliminary analysis of the mark-recapture data). At the population level we determined for each year a start, a peak, and an end date of breeding (i.e., first capture night, the capture night when most toads were captured, and last capture night, respectively). These calendar dates were all transformed into days of the year (where January 1st = 1), to facilitate modelling of long-term trends. These dates come with a degree of uncertainty, given the sampling done every 2–4 days and not daily. The date of start of the breeding comes with additional uncertainty as the first capture night is not always reflective of the same toad activity at the pond over the study period. We accounted for these sources of uncertainty in all following analyses, using simulated data on start and peak breeding dates.

### Climatic data

We obtained climatic data for the period 1980–2020 from the DaymetCH dataset (data obtained from *Bioclimatic maps of Switzerland © WSL, based on station data from the Federal Office of Meteorology and Climatology MeteoSwiss*, and elaborated by the *Land Change Science group, WSL*). This dataset consists of a 100-metre resolution grid of interpolated estimates of weather variables, using meteorological data from ground stations and the Daymet software (Thornton et al., 1997). We obtained data for the cell containing the breeding pond for the following variables: daily minimum, maximum, and mean temperature, daily total precipitation, and daily snow water equivalent (SWE; the equivalent amount of water stored in the snowpack). We then calculated average seasonal minimum, maximum, and mean daily temperatures, and cumulative seasonal precipitation and SWE.

### Data analysis

#### Population trend

A visual inspection of the data suggests that the trends in the breeding phenology across the study period are non-linear, both for start and peak breeding (Figure 1). Therefore, to better describe the observed trends, we conducted a piecewise regression on both start and peak breeding using the R package *segmented* (Muggeo, 2008). This analysis enables the identification of possible breakpoints in a trend, in our case a year (or several years) when a significant change occurs in the temporal trends of the breeding phenology. We set the year 1982 as year 0 in the model, to obtain a more intuitive interpretation of the intercept. Moreover, we decided to assess the robustness of our analysis to possible imperfect assignment of start and peak dates, as the toad sampling is not done daily. To do this, we simulated 1000 datasets of breeding start dates over the study period, allowing the date of the start of the breeding to be as early as seven days before the originally assigned first capture night. The process was described by a uniform distribution, where each date between 0 and 7 days earlier than the assigned date had the same probability of being chosen. We also simulated 1000 datasets for peak breeding dates, allowing the dates to deviate from the originally assigned date by letting it vary between the previous and the following capture night, again with the dates being picked from an uniform distribution. Using these simulated datasets, we ran 1000 piecewise regressions for both start and peak breeding dates, and calculated the 2.5^th^ and the 97.5^th^ percentiles of the values of each model parameter, including p-values testing for the significance of the breakpoint.

**Figure 1.**
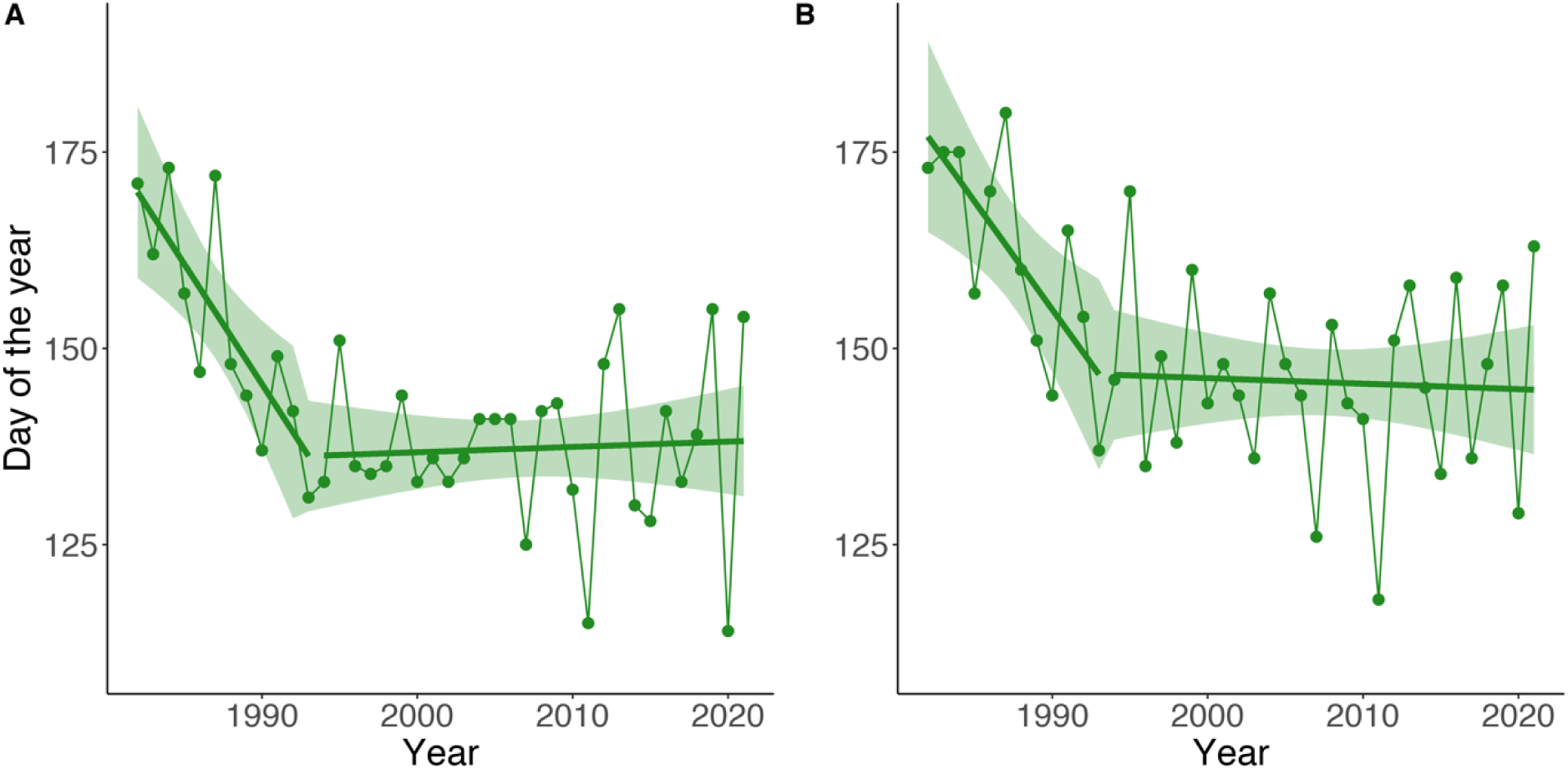
Trends of breeding phenology over the study period (1982–2021). **(A)** First day of the breeding season (day of the year, where January 1st = 1). The segmented green line is the result of a piecewise regression, where the year 1993 (± 5; 95% CI) was identified as a breakpoint, thus creating two distinct trends. **(B)** Date of peak breeding (i.e., date where most toads were captured in a given breeding season. The segmented green line is the result of a piecewise regression, where the year 1993 (± 6, 95% CI) was identified as a breakpoint. The green band in both plots represents the 95% CI for the piecewise regression.

Moreover, to check how the standard deviation (SD) of the start or the peak breeding dates changes over time, we calculated for both start and peak breeding the SD of the residuals of each of the 1000 piecewise regressions, using a rolling window approach (with a 10-year window) with the function *rollapply* of the package *zoo* (Zeileis & Grothendieck, 2005).

#### Determinants of variation in the breeding phenology in the population

To understand the climatic causes of the observed shifts in the breeding phenology of this population, we investigated the effects of several climatic variables on the timing of breeding at the population level. We identified *a priori* the climatic covariates that most reasonably could influence the breeding phenology in spring based on previous literature and expert knowledge (Oseen & Wassersug, 2002; Reading, 2003; While & Uller, 2014; Ficetola & Maiorano, 2016; Green, 2017). These climatic covariates are: average minimum daily temperature in spring (TSp) and winter (TW), total precipitation in spring (PrecSp, which includes both rainfall and snowfall), total snow water equivalent in spring (SWESp), and winter (SWEW). We then performed a piecewise regression on the time series of these five climatic covariates (Figure 2, Table S4). We used minimum temperatures because toads are nocturnal animals and are therefore more exposed to colder temperatures and less to average or warmer temperatures. Moreover, minimum temperatures will determine if the ground stays above freezing conditions. Changing the temperature variable (mean vs minimum vs maximum) in the subsequent analyses did not change the results as they were highly correlated (r > 0.93).

**Figure 2.**
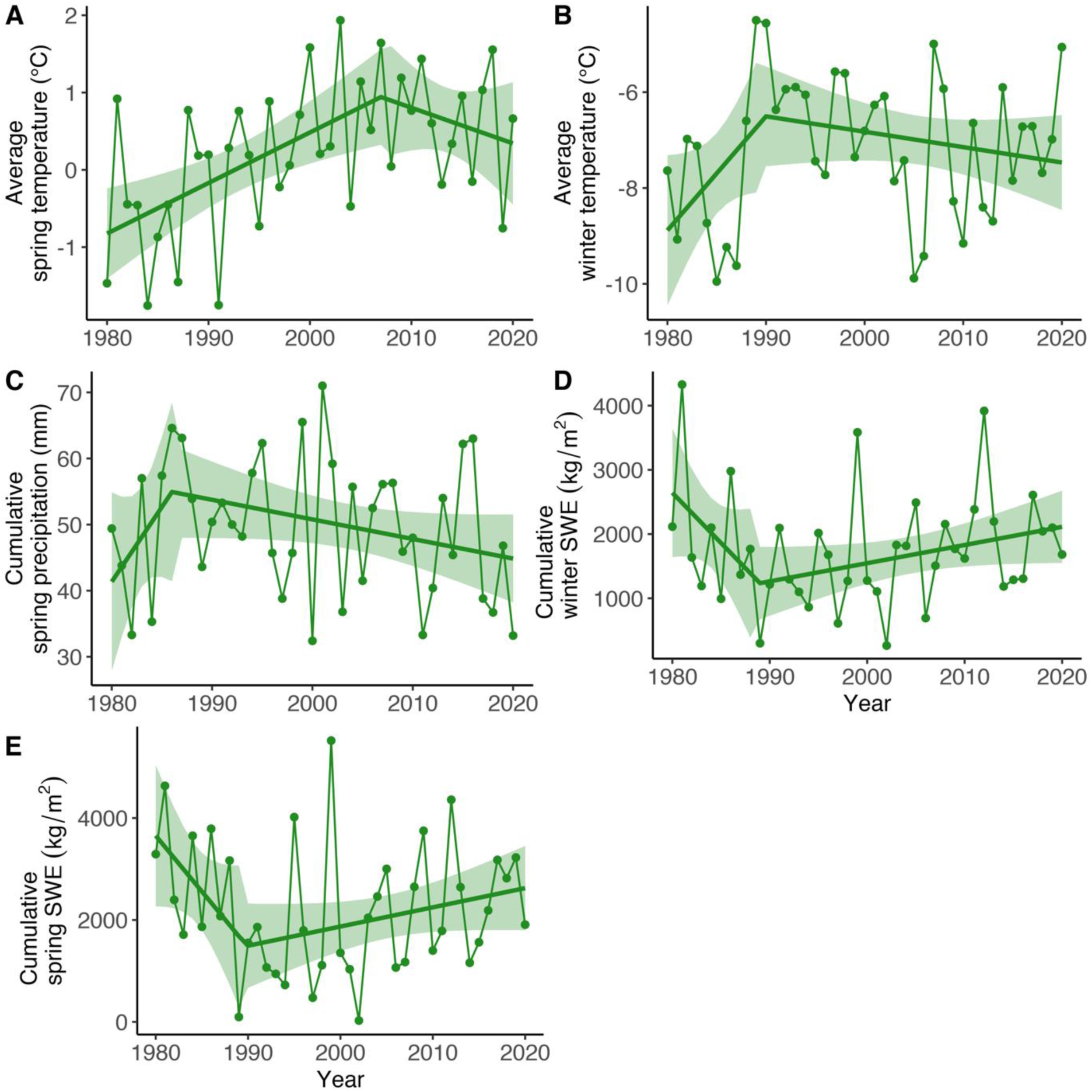
Trends over the study period of the five focal environmental variables. **(A)** Average minimum daily temperature in spring. The piecewise regression identified the year 2007 (± 9, 95% CI) as a breakpoint. **(B)** Average minimum daily temperature in winter. The year 1990 (± 7, 95% CI) was a breakpoint. **(C)** Cumulative precipitation in spring. The year 1986 (± 8, 95% CI) was a breakpoint. **(D)** Cumulative snow water equivalent (SWE) in winter. The year 1989 (± 7, 95% CI) was a breakpoint. **(E)** Cumulative SWE in Spring. The year 1990 (± 7, 95% CI) was a breakpoint. In all plots green ribbons represent the 95% CI for the linear regressions. Table S4 in the Appendix shows the summary of these five piecewise regressions.

With warmer winters and springs, toads should emerge sooner from their hibernation burrows as the snow will melt and the ground unfreeze earlier (Corn, 2003). The higher the snow water equivalent, the later the toads will emerge, as the snow cover will keep them blocked underground (Corn, 2003). Finally, precipitation can either favour or delay the breeding season. Snowfall should delay breeding as the snow cover will increase (Corn, 2003), but rainfall could potentially lead to an earlier start of the breeding season, as toads need high humidity levels to be active (Todd et al., 2011; Green, 2017). After standardising these climatic variables by subtracting the mean value and dividing by the standard deviation, we performed a principal component analysis (PCA, function *prcomp*, R package *stats* (R Core Team, 2020)), to reduce dimensionality and obtain uncorrelated variables (Figure S1).

In addition to these five climatic variables, the lunar cycle has also been identified to be an important factor for the timing of breeding in amphibians, with in general peak migration to the breeding site under waxing or full moon phases (Grant et al., 2009; Arnfield et al., 2012; Green et al., 2016; Jarvis et al., 2021). To assess the effect of the lunar cycle on the breeding phenology in our population, we first obtained the moon phase for each date of breeding start and peak breeding over the study period using the package *lunar* (Lazaridis, 2014). Following Arnfield et al. (2012) and Jarvis et al. (2021), we transformed the lunar phases in lunar angles (in radians, where 0 = new moon and π = full moon).

To quantify both the effects of climate and of the moon cycle on the breeding phenology, we modelled two separate linear regressions on the day of the breeding start and the day of peak breeding over the period 1982–2020. As explanatory variables we used the scores of the first two principal components (PC), as they explained an important amount of the variance in the data (>70%). As an additional explanatory variable, to better understand the role of the moon cycle, we included the cosine of the lunar angles of the start and peak breeding dates respectively. We first modelled the originally assigned dates, and then, as we did for the piecewise regression, we ran 1000 models with simulated datasets with varying dates of start and peak breeding, drawn from an uniform distribution. Each date could vary to be any date between the previous and following capture night.

To further study the association between the moon cycle and breeding phenology we tested if start and peak breeding tended to happen more frequently under certain moon phases. To do this, we used the *rayleigh.test* function of the *circular* R package (Agostinelli & Lund, 2017) to perform the Rayleigh test, a circular goodness-of-fit test that is particularly suited for checking if the values of a circular variable show a unimodal departure from a uniform distribution (Landler et al., 2018). To check for significant multimodal departures we performed the Hermans-Rasson test instead, using the *HR_test* from the *CircMLE* package (Fitak & Johnsen, 2017; Landler et al., 2018). Both tests were performed on the values in radians of the lunar angles. Also in this case we first ran the tests on the originally assigned dates and then we ran them on 1000 simulated datasets of start and peak breeding dates and obtained the 2.5^th^ and 97.5^th^ percentile of the p-values.

#### Determinants of individual variation in breeding phenology

In addition to considering phenology at the population level, we also wanted to understand whether individuals can show different patterns of changes in their reproductive phenology over time through different responses to climatic variables, possibly indicating a genetic component that mediates the effect of the changing environment. We therefore modelled the effect of the previously used principal components PC1 and PC2, as well as of the cosine of the lunar angle on each individual first capture occasion in any given year (6735 occurrences for 3053 uniquely marked individuals, as many individuals were breeding in multiple years (mean = 2.21 years, SE = 0.02)), using a linear mixed model (package *lmerTest*; Kuznetsova et al., 2017). Also in this case, we first ran the model on the originally assigned arrival dates, and then, to account for uncertainty in the assignment of the dates of arrival to the pond we simulated 1000 new datasets where every individual arrival date is newly sampled from an uniform distribution and can be as early as the capture night preceding the original arrival date, or if it was the first capture night of the season, up to seven days before. Using these 1000 new datasets we ran 1000 models and obtained the 2.5^th^ and the 97.5^th^ percentile values for each parameter. As a random effect, applied on both the intercept and the slope of both PC1 and PC2, we included individual identity (ID). This was done not only to observe if individuals react differently to changing environmental conditions, but also to account for the non-independence of the data. Moreover, we also included *year* as a random effect on the intercept, to account for unexplained year-specific variation in the data. Finally, we included the effect of sex to account for differences between males and female. To properly be able to compare the effects of continuous variables (i.e., the two PCs and the cosine of the lunar angles) with the effect of a categorical variable (i.e., sex), we standardised the three continuous variables by subtracting the mean and dividing by two times the standard deviation (Gelman, 2008). Finally, as a measure of model fit, we calculated the conditional R^2^ value using the *r.squaredGLMM* function from the package *MuMIn* (Barton, 2019).

#### Repeatability of arrival date

Finally, we also estimated repeatability (i.e., the upper limit of heritability) of arrival dates at the breeding site. High values of repeatability (*r*) mean that individuals are consistent in their relative arrival timing (e.g., always among the first ones), and vice versa. To calculate *r*, we used for each individual the date of first capture for each year that it was captured. This date is a relatively good proxy for the date of arrival at the breeding site, as the data collection usually starts every year approximately when the first toads arrive at the pond. The date was converted to the day of the year (where January 1st = 1), and then standardised by subtracting the year-specific mean and dividing by the year-specific standard deviation. We then used the function *rpt* from the package *rptR* to calculate *r* using individual ID as the group variable (Stoffel et al., 2017), and bootstrapping 1000 times to obtain the 95% CI. As for all the other analyses, to account for the uncertainty in the assignment of the dates, we repeated the calculation of *r* 1000 times, sampling different arrival dates every time from a uniform distribution, where the arrival date of each individual can be up to the previous capture night, or up to seven days earlier if they were caught during the first capture night of the season. We then calculated the 2.5^th^ and the 97.5^th^ percentiles of *r* to show the spread it can have. Given the different reproductive strategies that males and females toads have, with females on average coming to the breeding site later than males and for a shorter period of time (Reading & Clarke, 1983; Loman & Madsen, 1986), we performed sex-specific calculations of *r*.

We conducted all the analyses in R (R version 4.1.1; R Core Team, 2020) with RStudio (version 2022.7.1.554; R Studio Team, 2022).

## Results

### Population trend

Both the breeding start dates and the dates of peak breeding show very similar trends (Pearson’s correlation coefficient = 0.91), with both also showing marked between-year variation over the study period. Nonetheless, a shift towards earlier breeding dates is observable, with breeding happening now on average around 30 days earlier compared to the start of the study period (Figure 1). The piecewise regression on breeding start dates identified a single breakpoint in the temporal trend in the year 1993 with a pre-1993 steep advancement of breeding dates followed by a post-1993 almost flat trend (Figure 1A; Table 1). The analysis of the robustness of the piecewise regression, done by simulating data and running 1000 piecewise regressions, performed very similarly, with 910 cases out of 1000 where the year 1993 was identified as breakpoint and the model coefficients were very close to the piecewise regression conducted on the originally assigned breeding dates (Table S1). The piecewise regression on peak breeding dates also identified 1993 as a breakpoint year (Figure 1B; Table 1). In this case, the analysis of the robustness showed slightly more variation, with the breakpoint years mostly obtained being 1993 and 1996 (274 and 283 out of 1000 respectively) (Table S1). Moreover, we found the standard deviation (SD) of the residuals of the piecewise regressions on both start and peak breeding dates to vary considerably, with higher SDs at the start and the end of the study period (Figure S2). To further check the pattern in the residuals we split them in four different decades and checked their distribution (Figure S3).

**Table 1.**
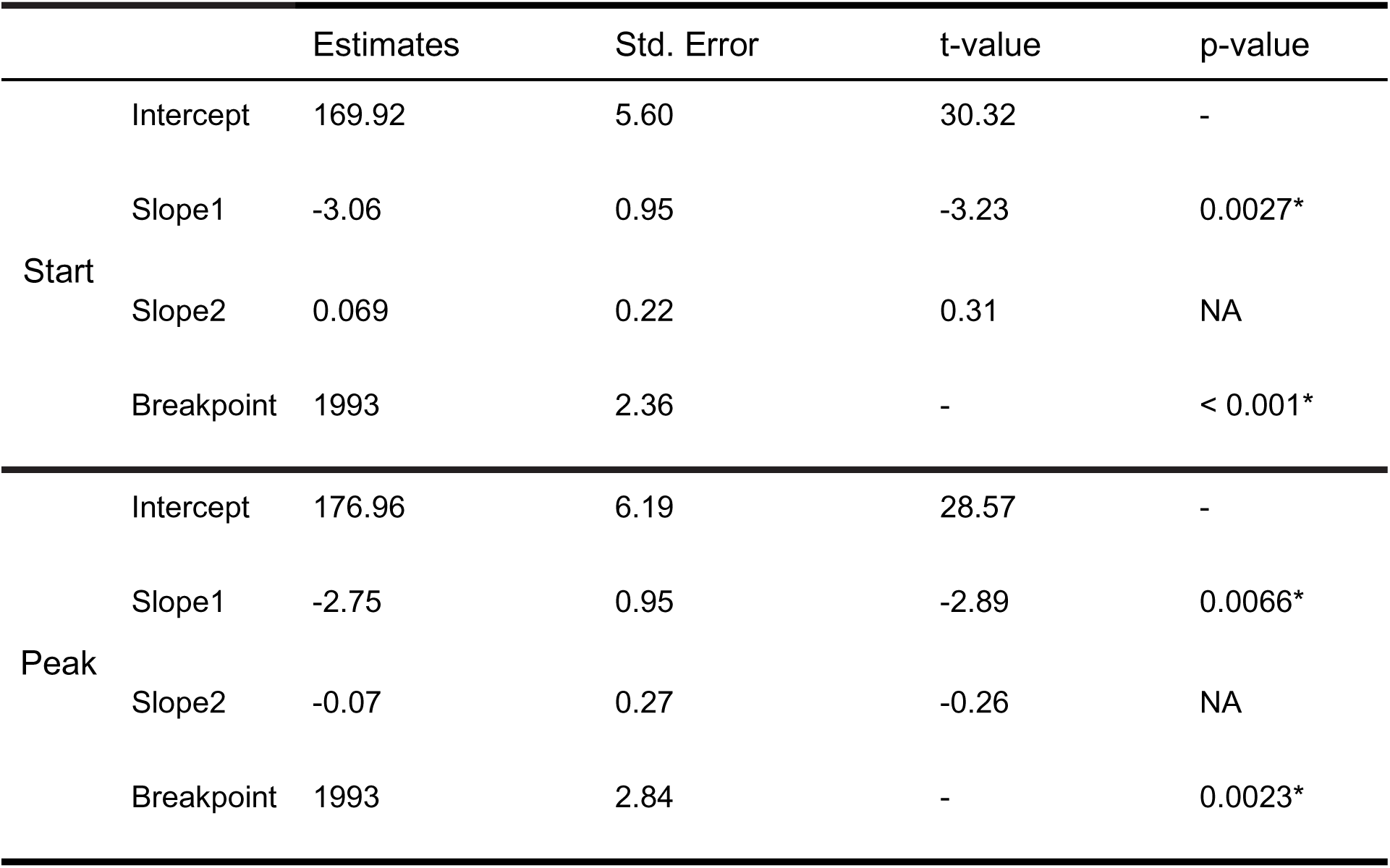
Summary of the piecewise regressions on the start and peak of the breeding season. For both intercept and slopes we show the estimate, its standard error, and the t-value and p-value associated with it. Slope1 refers to the segment before the breakpoint and Slope2 refers to the segment after the breakpoint. Asterisks next to the p-values show significance at the 0.05 level. The p-value for Slope2 is NA since standard asymptotics do not apply (Muggeo, 2008). No p-values are provided for the intercept because this test is not of biological interest.

### Determinants of variation in the breeding phenology in the population

The first two principal components (PC) of the principal component analysis (PCA) described together more than 70% of the variation in the data, and both had a standard deviation (i.e. the squared root of their eigenvalue) above one (Figure S1; Table S5). Therefore, applying the Kaiser rule, we kept the scores of these two PCs (PC1 and PC2) as explanatory variables in the following linear regressions on the start of the breeding season and on peak breeding (also including the scaled cosine of lunar angle). PC1 was mostly determined by winter temperature (+0.45 loading) and winter and spring SWE (-0.61 and -0.64, respectively). PC2 was mostly determined by spring weather conditions. Spring temperature had a negative loading (-0.68), while precipitation had a positive loading (+0.68) (Figure S1; Table S6).

Regarding the start of the breeding season, the model (adjusted *R^2^* = 0.41) indicated a significant negative relationship with PC1 and a significant positive relationship with PC2 (Table 2). The cosine of the lunar angle had a non-significant effect. Similarly, for the regression on the dates of peak breeding, we found a significant negative relationship with PC1 and a significant positive relationship with PC2, while the cosine of the lunar angle had a small and non-significant effect (Table 2). The adjusted *R^2^* was 0.54. In both cases the outcome is that warmer temperatures in winter and spring, less snow cover, and weaker precipitations are all associated with an earlier start and peak of the breeding season. Both the 1000 linear regressions on the simulated dates of the start of the breeding season and the 1000 on the simulated dates of peak breeding performed similarly to the two regressions on the originally assigned dates (Table S2), indicating that our analysis is robust to possible imperfect assignment of dates of start and peak breeding.

**Table 2.** *Summary of the linear regression on the start and peak of the breeding season. For each variable we show the estimate, its standard error, and the t-value and p-value associated with it. Asterisks next to the p-value show significance at the 0.05 level. No p-values are provided for the intercept because this test is not of biological interest*.

|  |  | Estimates | Std. Error | t-value | p-value |
| --- | --- | --- | --- | --- | --- |
| Start | Intercept | 141.72 | 1.65 | 85.92 | - |
|  | PC1 | -5.48 | 1.67 | -3.28 | 0.0024* |
|  | PC2 | 7.00 | 1.68 | 4.17 | 0.00019 |
|  | cos(moon) | 1.32 | 1.68 | 0.79 | 0.44 |
| Peak | Intercept | 150.21 | 1.56 | 96.59 | - |
|  | PC1 | -5.47 | 1.58 | -3.47 | 0.0014* |
|  | PC2 | 9.37 | 1.60 | 5.86 | < 0.0001* |
|  | cos(moon) | 0.24 | 1.60 | 0.15 | 0.88 |

### Effect of the moon cycle on breeding phenology

To further understand if the lunar cycle is associated with the breeding phenology, we performed two statistical tests. To check for unimodal deviation we ran a Rayleigh’s test on the moon phases on breeding season start and on peak dates. In both cases we obtained a non-significant p-value (0.27 and 0.08 respectively), indicating that we could not confidently reject the null-hypothesis of the data being uniformly distributed in the circular space. In addition, the outcome of the Hermans-Rasson test for multivariate deviations indicated that the null hypothesis could not be rejected for both start and peak breeding (p-value = 0.38 and 0.21 respectively). To further assess the robustness of our analysis to imperfect assignment of dates we ran both the Rayleigh’s and Hermans-Rasson test on 1000 simulated datasets of dates of start and peak breeding. The outcome is similar to the tests performed on the originally assigned dates. The p-values of the Rayleigh’s test were 0.36 [2.5^th^ and 97.5^th^ percentiles: 0.07; 0.80] and 0.16 [0.008; 0.60] respectively. The p-values for the Hermans-Rasson test on start and peak breeding were 0.43 [0.05; 0.90] and 0.25 [0.011; 0.78] respectively. This means that there was no clear pattern between lunar phases and the start of the breeding season or the peak breeding (Figure 3).

**Figure 3.**
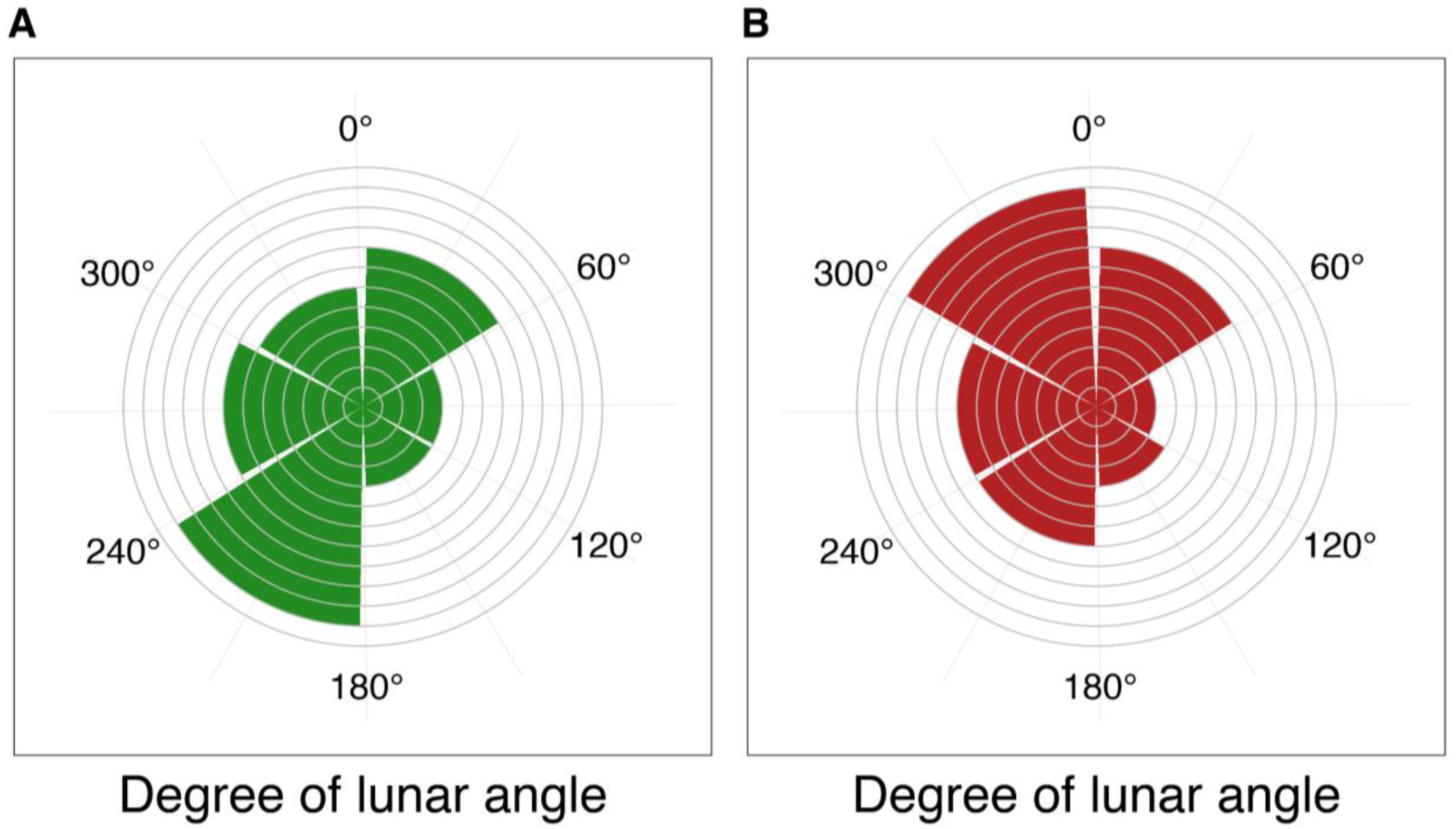
Circular histogram showing counts of **(A)** the originally assigned breeding start dates and **(B)** the originally assigned peak breeding dates under different lunar phases for the period 1982–2021 (e.g., the breeding season started eight times under a moon phase with a lunar angle between 0° and 60°). Lunar angles, initially in radians, were back-transformed to degrees, so that the new moon is at 0° and full moon is at 180°.

### Determinants of individual variation in breeding phenology

To better understand if there are among-individual differences in the phenological response to changing climatic variables, we used a linear mixed model to test for the effect of climatic variables on the individual breeding start dates (i.e., the date on which an individual was first captured). We found only a small difference in the response of breeding phenology to climatic variables among individuals (i.e., low values for the random effect ID, both on intercept and slopes, Table 3). We found a strong significant positive effect of PC2 on the breeding dates (17.51 ± 3.27 SE), meaning that stronger precipitation and lower minimum spring temperatures are associated with a delay in the breeding. We also found a significant and strong negative effect of PC1 (-10.14 ± 2.85 SE), indicating that colder winter temperatures and higher SWE are associated with a delay in the breeding. We also found a significant but weak effect of the cosine of the lunar angle (1.57 ± 0.14 SE), suggesting a possible small role of the lunar cycle. Finally, we observed an effect of sex indicating that males arrived on average earlier than females (-1.45 ± 0.14 SE) (Table 3). The 1000 models on the 1000 simulated datasets, ran to assess the robustness of the analysis to imperfect assignment of arrival dates, showed a similar outcome to the main model (Table S3).

**Table 3.** *Detailed description of the model used to check for the effect of environmental variables on the phenology at the individual level. Sex is included to observe differences between males and females. The response variable ArrivalDate is a vector of dates of arrival at the breeding site for each individual over the study period. PC1 and PC2 are the first two components of the PCA performed on the climatic data. Cos(moon) is the cosine of the lunar angle for the arrival date. ID refers to the identity of each individual, and it is used as a random effect on both intercept and the slopes of PC1 and PC2. Finally, Year is included as a random effect to account for additional unexplained variation that might be caused by sampling variation. The second and third part of the table provide details on the estimates for the fixed and random effects respectively. No p-values are provided for the intercept because this test is not of biological interest*.

| Model name | Variables |  |  |  | Conditional R<sup>2</sup> |
| --- | --- | --- | --- | --- | --- |
| Full_model | ArrivalDate ~ PC1 + PC2 + Sex + cos(moon)<br>+ (1\|Year) + (1\|ID) + (0 + PC1\|ID) + (0 + PC2\|ID) |  |  |  | 0.92 |
|  | Effect size | Std. Error | t-value | P-value |  |
| Intercept | 147.87 | 1.66 | 89.24 | - |  |
| Sex (male) | -1.45 | 0.14 | -10.44 | < 0.0001* |  |
| PC1 | -10.14 | 2.85 | -3.56 | 0.0011* |  |
| PC2 | 17.51 | 3.27 | 5.36 | < 0.0001* |  |
| cos(moon) | 1.57 | 0.14 | 11.08 | < 0.0001* |  |
| Variance |  |  |  |  |  |
| ID (intercept) | 2.47 |  |  |  |  |
| ID on PC1 | 2.89 |  |  |  |  |
| ID on PC2 | 0.58 |  |  |  |  |
| Year (intercept) | 101.78 |  |  |  |  |
| Residuals | 17.73 |  |  |  |  |

### Repeatability of arrival date

In total, 453 females and 1092 males visited the pond over multiple years. The repeatability value calculated with the originally assigned arrival dates was 0.15 [95% CI 0.08; 0.21] for females and 0.12 [95% CI 0.09; 0.15] for males. To again assess the robustness of our analysis we simulated 1000 new datasets with varying arrival dates and calculated 1000 repeatability values for females and 1000 for males. We found a mean repeatability value *r* of 0.14 [2.5^th^ and 97.5^th^ percentiles 0.12; 0.17] for females and 0.10 [2.5^th^ and 97.5^th^ percentiles 0.09; 0.11] for males.

## Discussion

Our results show that variation in the breeding phenology is strongly associated with climatic conditions, which vary substantially among years but also show trends across times. We also found low repeatability values and low variability in individual responses, suggesting that the genetic component contributing to the observed variation of individuals in the breeding phenology is weak. Finally, we found indications of a possibly significant, but weak, effect of the lunar cycle. A signal might indeed exist, but the climatic variables probably have a stronger effect.

Our results support the hypothesis of a strong link between the breeding phenology of high-elevation amphibian populations and climatic conditions. Increasing temperatures are a key driver of snow melt and ground defrosting, which in turn act as important environmental cues for toads to initiate migration to their breeding grounds (Corn & Muths, 2002; Green, 2017). During particularly warm springs, the snow melts and the ground defrosts earlier, leading to a shift of the onset of breeding to earlier dates. Our findings on the importance of temperature are in line with previous studies on *B. bufo* (Reading & Clarke, 1983; Reading, 2003; Tryjanowski et al., 2003; Arnfield et al., 2012). On the other hand, where past studies have identified rainfall to be an important trigger for migration in lowland populations (Reading & Clarke, 1983; Sinsch, 1988; Jarvis et al., 2021), we did not clearly observe this in our data, as our measure of precipitation included both snow- and rainfall. We found that a higher amount of precipitation in spring (combined with a decrease of spring temperature) was associated with a later breeding date. In fact, at low temperatures, precipitation in the form of snowfall or freezing rain can delay the melting of the snow cover, therefore leading to a delay in the breeding. The observed negative association between snow water equivalent (SWE) and breeding timing is in line with the rest of the findings. In fact, SWE depends considerably on temperatures and precipitation, as well as other aspects such as exposition, and it is a key factor that influences phenology (Corn, 2003). The very similar trend observed for peak activity in breeding indicates that both start and peak breeding are influenced mostly in the same way by the same climatic variables.

When looking at the individual timing of arrival we still found an important effect on the breeding phenology of PC1 (TW and SWESp/W) and PC2 (TSp and PrecSp) (Table 3). However, we found only non-significant and small among-individual variation in phenological response to changing climatic conditions (Table 3). As reproduction happens only once a year in explosive breeders living in temperate zones, synchronisation in breeding could be key to maximise reproductive output (Ims, 1990). Such an accurate synchronisation can be achieved more easily when all individuals hibernating close to each other express similar responses to external cues triggering their migration to the breeding pond, instead of responding individually in different ways, highlighting once more that the breeding phenology is mainly driven by climatic conditions.

Moreover, the low values of *r* (i.e., the upper limit of heritability) that we found for the timing of arrival show that there is some individual heterogeneity in this trait, and it could further indicate that there is only a small contribution of the genetic component to variation in the breeding phenology. This conclusion is in line with what most studies on amphibian phenology found (Semlitsch et al., 1993; Blaustein et al., 2001; Parmesan, 2007; While & Uller, 2014; but see Heusser & Ott, 1968; Phillimore et al., 2010). In other species, for instance birds, higher values of repeatability have been found for migration phenology, a trait linked to breeding. Franklin et al. (2022) found in their meta-analysis an average value of repeatability of 0.414, while Kürten et al. (2022) found repeatability values above 0.60 for various traits (but see Clermont et al., 2018; Vaillant et al., 2021 for examples of low repeatability in birds), but in amphibians that follow an explosive breeding strategy, the genetic component does not appear to be the main determinant of variation in breeding phenology. This might be due to either populations being truly able to respond plastically to changing climatic conditions, and therefore there is no strong selection on genetic variation in the trait, or there might be little genetic variation in the population to begin with. Low values of repeatability might also indicate a non-consistent choice of the hibernation site (and therefore distance to the pond). Not much is known about hibernation site fidelity in anurans, and future studies should address this question.

Finally, we found that on average males tend to arrive earlier than females (Table 3), similarly to what has been found in lowland populations of *B. bufo* (Loman & Madsen, 1986; Höglund & Robertson, 1987, 1988; but see Gittins et al., 1980). In these studies, males, especially bigger ones, were observed to arrive on average earlier at the breeding pond. Smaller males, on the other hand, were observed intercepting females on their way to the pond, betting on the fact that the females would lay the eggs as soon as they arrived at the pond, avoiding competition from the other bigger males. A more detailed future analysis of body size and its effects on the timing of migration to the breeding site could confirm this theory also for our study population.

Climate change is leading to on-average increasing temperatures both globally but also at smaller scales such as in the European Alps (Vitasse et al., 2021) and in Switzerland (Rebetez & Reinhard, 2008). The start of data collection for this study (early 1980s) coincides with an important increase of temperatures in Switzerland (Bundesamt für Umwelt (BAFU), 2020). In fact, each year since the mid-80s, the deviation from the mean yearly temperature (average calculated over the period 1864–2019) has always been positive (Begert & Frei, 2018). In the Swiss Alps, mean temperature increased by about 1.7 °C from 1975 to 2004, nearly twice the global average (Rebetez & Reinhard, 2008). Despite these general trends, we observe at our study site stable or even decreasing trends in temperatures during the study period, especially in the second half (Figure 2). Initially, the shift towards earlier breeding (pre-1993/1996) can be explained by warming temperatures and decreasing SWE (Figure 2). On the other hand, the absence of a trend in the breeding dates observed after the mid-1990s (Figure 1) could be explained by a change in trajectories of winter temperature, which started decreasing around 1990 (Figure 2), as well as of winter and spring SWE, which started increasing around the same time. These combined changes are acting against the increasing spring temperature (which has increased until around 2007; Figure 2), therefore slowing down and ultimately halting the shift towards earlier breeding dates of the toads.

While we could expect climate change to act linearly on the shift towards earlier breeding dates, it is possible that other site-specific conditions prevail at different temporal and geographical scales, creating an heterogenous mosaic of climate conditions. An example of this is the influence of the North Atlantic Oscillation (NAO) on the breeding phenology of amphibians and reptiles in southern France, where shifts in the breeding phenology in the last forty years were related to variation in the NAO index (Prodon et al., 2020). High elevation habitats can also show different climates at very small geographical scales (Scherrer & Körner, 2011; Feldmeier et al., 2020). The phenology of populations experiencing these different microclimates will therefore not necessarily be affected in the same way (Miller et al., 2018; Arietta et al., 2020; Turner & Maclean, 2022). In our case, the data on climatic variables was limited to the 100 metres x 100 metres cell which includes the pond, and since we do not exactly know where the toads hibernate in the surrounding landscape, we cannot exclude that they are experiencing different microclimates compared to the pond and its surrounding area. Hibernating toads have been found more than 1000 metres away from the breeding site horizontally, and up to almost 400 metres away vertically (Sztatecsny & Schabetsberger, 2005). Since the breeding pond and surrounding area are often still partially covered by snow during peak night, the hibernation sites are probably warmer than the breeding site itself. Differences in microclimates between hibernation sites and breeding site could further explain individual variation in breeding timing (e.g., arrival at the pond). Further studies on how the hibernation sites of the toads in this population can affect the breeding phenology should be conducted.

Despite the observed stabilisation of the trend of the breeding dates (Figure 1), the study population appears to experience increased variation in the dates of the start of the breeding season (Figure S2 and Figure S3). This increased variation could be explained by extreme weather events whose occurrence is expected to increase under climate change (Rahmstorf & Coumou, 2011; National Academies of Sciences, Engineering, and Medicine, 2016). Such unpredictability and extremeness of environmental conditions could threaten populations if they lead to either excessively early or late breeding, especially in temperate regions. In fact, extreme early breeding is associated with reduced hibernation periods which can decrease the body condition in spring (Reading, 2007). Additionally, early breeding can expose eggs and tadpoles to late frost events (Muir et al., 2014; Bison et al., 2021; Turner & Maclean, 2022). Delayed breeding can potentially pose a problem as well if the pond dries out during warm periods in late spring or if juveniles cannot accomplish full growth before hibernation. Indeed, smaller and younger juveniles are more at risk of death before and during the first hibernation period (Morin et al., 1990; Sinsch & Schäfer, 2016). This seems to be compensated in some cases by an accelerated growth at the larval stage in case of late breeding, but with a cost of reduced defences against predation (Orizaola et al., 2016). Such riskful situations can have strong negative effects on individual survival and reproductive output, ultimately leading to population declines (Reading, 2007; Iler et al., 2021). On the other hand, at least initially, climate change could lead to longer growing seasons during which individuals would have the opportunity to gather more energy before the onset of hibernation (Zani, 2008; Iler et al., 2021), with potentially positive effects at the population level. Climate change can as well lead to species expanding upward (Vitasse et al., 2021), with *Bufo bufo* populations observed locally extending their upper range limit to higher elevations (Lüscher et al., 2016). If moving upwards is not possible, high-elevation populations adapted to their environments could face local extirpation (Urban, 2018).

## Conclusion

In this study we showed the important association between climatic variables such as temperature, snow cover, and precipitation with the breeding phenology of a *Bufo bufo* population living at high elevations. Breeding happens on average around 30 days earlier now compared to four decades ago, and interestingly the shift towards earlier breeding dates has not been constant, but is better described by two different trends. After an initial steep advancement until the mid-90s, the trend stabilised. This is reflected in the trends of the time series of the focal climatic variables, which explain the observed temporal variation in breeding phenology. The stabilisation in the trend suggests that there might be spatial heterogeneity in climate change and its effects, therefore different populations might show different trends in their breeding phenology. This stabilisation is accompanied by an increased variation in the dates of the start of the breeding season, with potential consequences for the population that should be further investigated in the future. To conclude, this 40-year study is one of the first and most detailed studies on the breeding phenology of alpine populations of *B. bufo*, and it highlights the influence of changing environmental conditions on the timing of reproduction.

## Acknowledgements

We warmly thank all the people involved in data collection and management since 1982 and everybody who facilitated fieldwork during the past 40 years. Capture, handling and marking of toads were done under an animal welfare permit issued by the Veterinäramt des Kantons Bern. We also thank the Land Change Science of the Swiss Federal Research Institute WSL for kindly providing us with the climatic data. Finally, we thank Sergio Estay, Nigel Yoccoz and an anonymous reviewer for comments on a previous version of the manuscript.

## Authors contribution

O.L. and B.S. conceived the study. K.G., S.Z., S.A., B.L., D.S., M.T., and H.R. collected data. O.L. prepared and analysed the data. B.S and A.O. provided feedback on the analyses. O.L. wrote the paper with input from all authors.

## Data and script accessibility

Data and scripts for this publication are available on the Zenodo Repository: https://doi.org/10.5281/zenodo.7333319.

## Supplementary material

Extra tables and figures are available in the Appendix.

## Conflict of interest and disclosure

The authors of this preprint declare that they have no financial conflict of interest with the content of this article.

## Funding

This project was funded by the Federal Office for the Environment (contract numbers: 20.0001.PJ/46DBED0F1 and 06.0126-PZ Artenförderung / N494-1379) and the Stiftung Temperatio.

## Appendix

**Figure S1.**
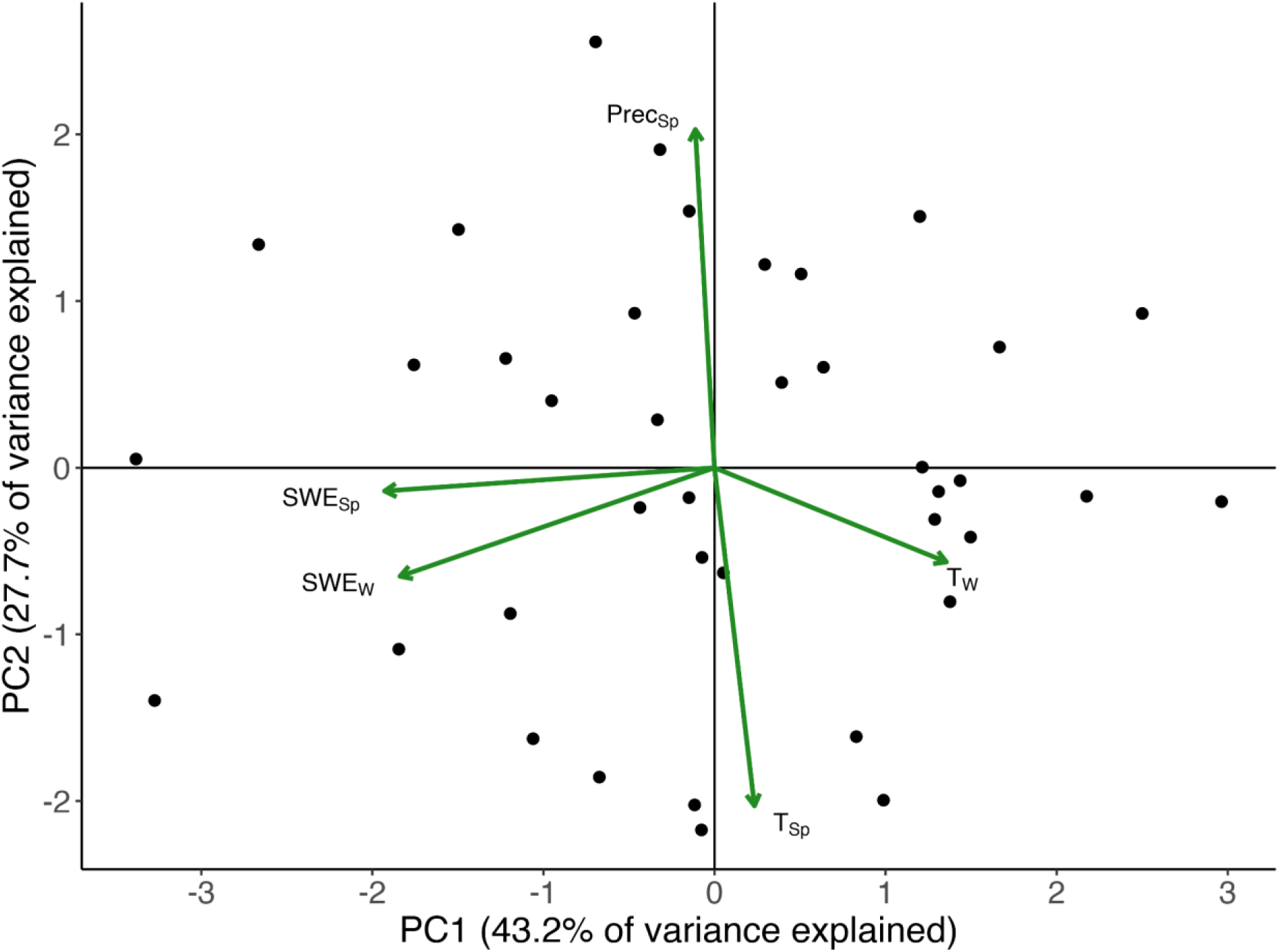
Graphical visualisation of the first two principal components of our principal component analysis. The points represent the scores over the two axes, while the arrows represent the loadings of the five environmental variables. The first principal component explained 43.2% of the variance in the data, while the second principal component explained 27.7%. TSp is the average minimum daily spring temperature, TW is the average minimum daily winter temperature, PrecSp is the total precipitation in spring, SWESp is the total spring snow water equivalent and SWEW is the total winter snow water equivalent

**Figure S2.**
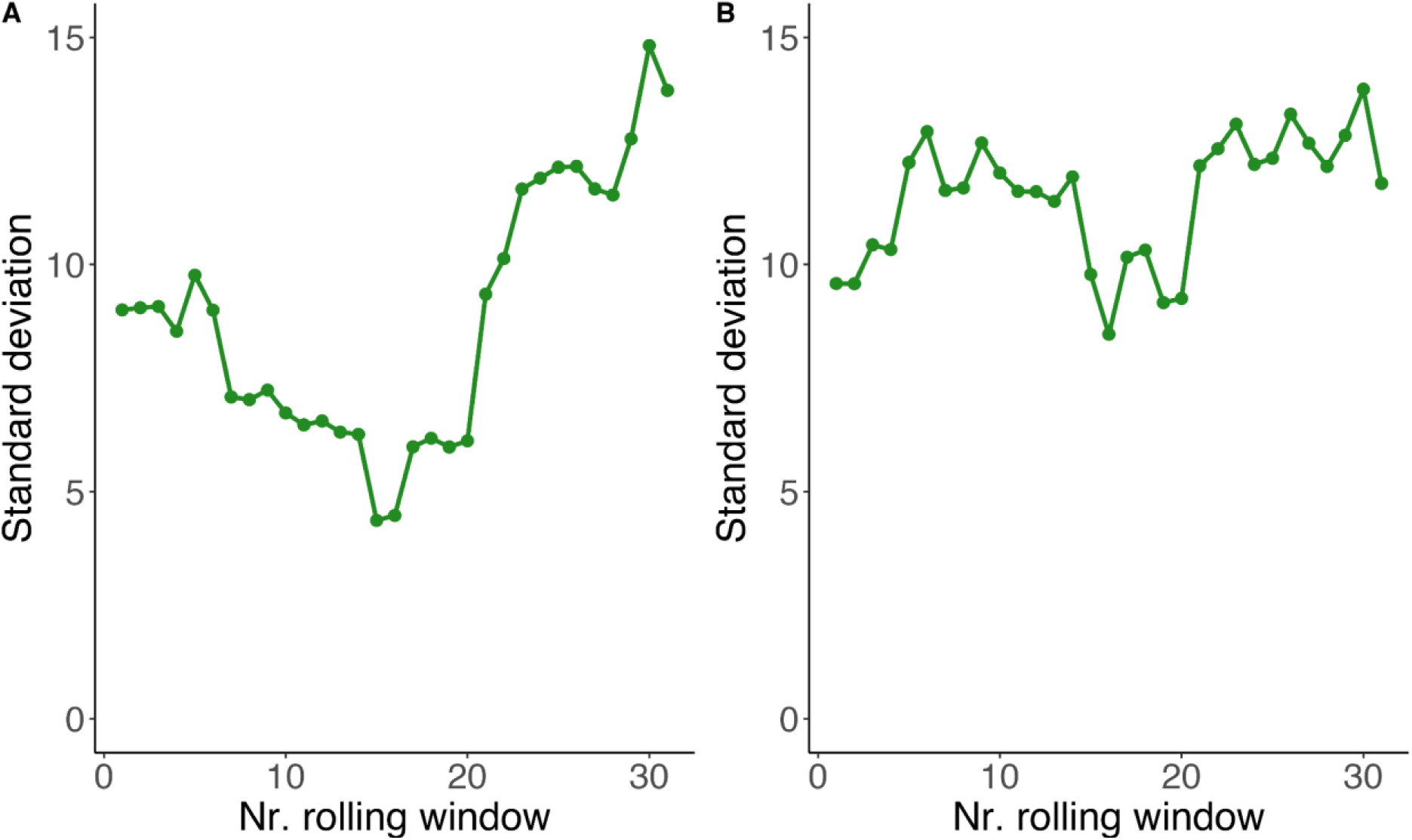
Mean standard deviation of the residuals of the 1000 piecewise regressions on start of the breeding season **(A)** and 1000 piecewise regression on peak of the breeding season **(B)**, calculated with a rolling window approach. Window size was 25% of the length of the timeseries (i.e., 10 years out of 40 years, therefore we had 31 windows).

**Figure S3.**
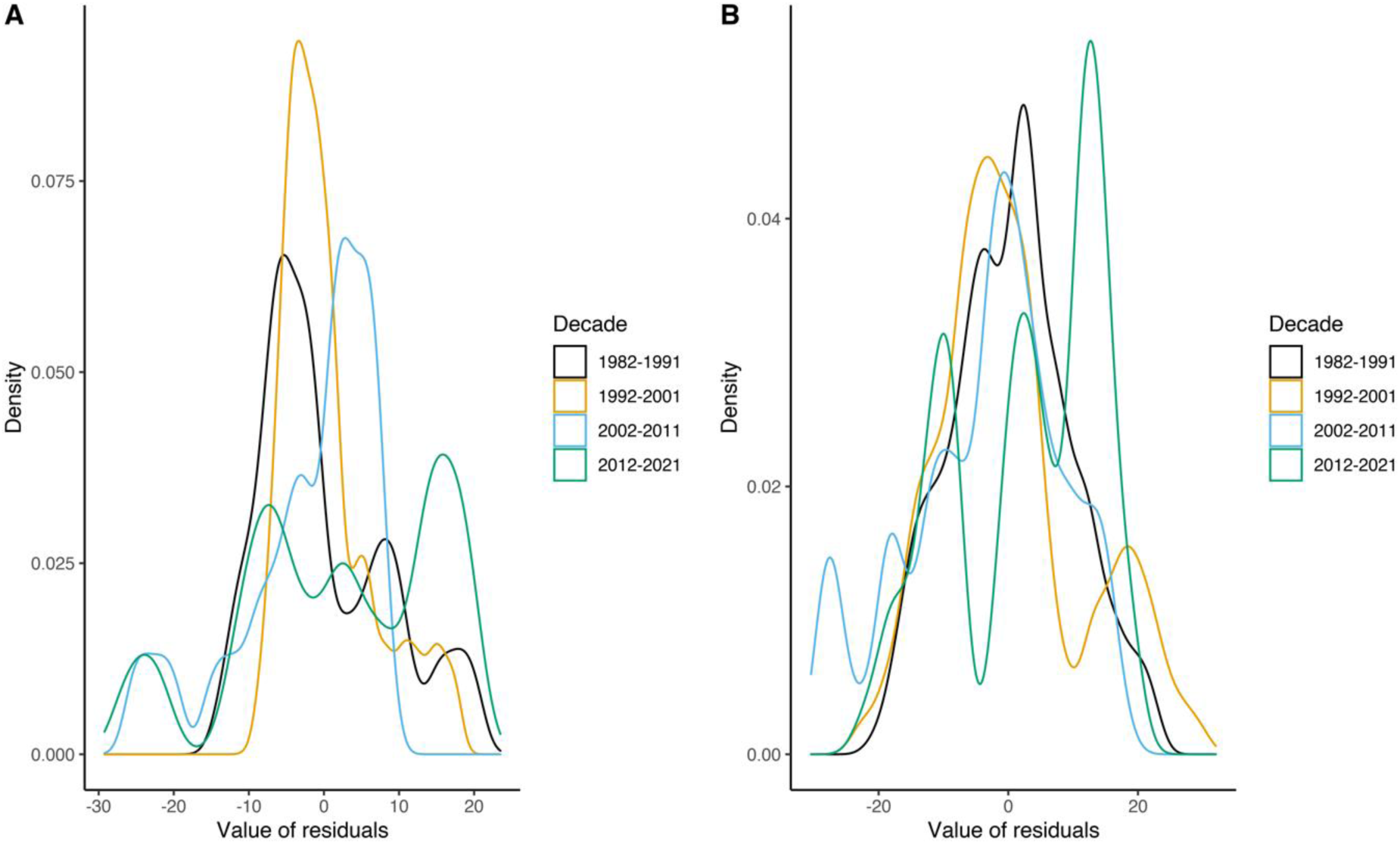
Distribution of the residuals of the 1000 piecewise regression on start of the breeding season **(A)** and peak breeding **(B)**. We divided the residuals in four different decades: decade 1 (1982-1991), decade 2 (1992-2001), decade 3 (2002-2011) and decade 4 (2012-2021).

**Table S1.** Summary of the simulated piecewise regressions on the start and peak of the breeding season. For both intercept and slopes we show the estimate, its standard error, and the t-value and p-value associated with it. Moreover, for each parameter we show in square brackets the 2.5th and the 97.5th percentiles of the values obtained by running 1000 models. Slope1 refers to the segment before the breakpoint and Slope2 refers to the segment after the breakpoint. Asterisks next to the p-values show significance at the 0.05 level. The p-value for Slope2 is NA since standard asymptotics do not apply (Muggeo, 2008). No p-values are provided for the intercept because this test is not of biological interest.

|  |  | Estimates | Std. Error | t-value | p-value |
| --- | --- | --- | --- | --- | --- |
| Start | Intercept | 166.59<br>[164.24 ; 169.07] | 5.70<br>[5.22 ; 6.17] | 29.29<br>[27.03 ; 31.87] | - |
|  | Slope1 | -3.11<br>[-3.60 ; -2.78] | 0.95<br>[0.81 ; 1.11] | -3.30<br>[-3.85 ; -2.86] | 0.0027*<br>[0.0005 ; 0.0069] |
|  | Slope2 | 0.062<br>[-0.043 ; 0.16] | 0.23<br>[0.21 ; 0.25] | 0.271<br>[-0.201 ; 0.679] | NA |
|  | Breakpoint | 1993 | 2.30<br>[1.93 ; 2.69] | - | < 0.001*<br>[0.00013 ; 0.0020] |
| Peak | Intercept | 174.47<br>[167.99 ; 179.56] | 5.52<br>[4.11 ; 6.88] | 32.34<br>[25.78 ; 41.35] | - |
|  | Slope1 | -2.10<br>[-3.10 ; -1.12] | 0.67<br>[0.26 ; 1.16] | -3.43<br>[-4.69 ; -2.38] | 0.0046*<br>[< 0.001 ; 0.022] |
|  | Slope2 | 0.41<br>[-0.11 ; 1.62] | 0.47<br>[0.25 ; 1.11] | 0.54<br>[-0.40 ; 1.64] | NA |
|  | Breakpoint | 1993 / 1996 | 3.17<br>[2.52 ; 3.83] | - | 0.0023*<br>[0.0005 ; 0.0064] |

**Table S2.** Summary of the simulated linear regressions on the start and peak of the breeding season. For each variable we show the estimate, its standard error, and the t-value and p-value associated with it. In square brackets we show the 2.5th and the 97.5th percentile values, obtained by simulating 1000 times the dates and running 1000 models. Asterisks next to the p-value show significance at the 0.05 level. No p-values are provided for the intercept because this test is not of biological interest.

|  |  | Estimates | Std. Error | t-value | p-value |
| --- | --- | --- | --- | --- | --- |
| Start | Intercept | 138.22<br>[137.54 ; 139.00] | 1.69<br>[1.58 ; 1.80] | 81.93<br>[76.79 ; 87.63] | - |
|  | PC1 | -5.52<br>[-6.34 ; -4.59] | 1.73<br>[1.60 ; 1.86] | -3.20<br>[-3.73 ; -2.60] | 0.0038*<br>[0.00068 ; 0.013] |
|  | PC2 | 6.99<br>[5.98 ; 7.84] | 1.75<br>[1.62 ; 1.90] | 4.01<br>[3.35 ; 4.69] | 0.00048*<br>[< 0.0001* ; 0.0019] |
|  | cos(moon) | 0.93<br>[-1.34 ; 2.86] | 1.77<br>[1.63 ; 1.93] | 0.53<br>[-0.72 ; 1.60] | 0.55<br>[0.11 ; 0.97] |
| Peak | Intercept | 150.35<br>[149.41 ; 151.28] | 1.58<br>[1.45 ; 1.70] | 95.43<br>[88.14 ; 104.06] | - |
|  | PC1 | -5.37<br>[-6.42 ; -4.32] | 1.61<br>[1.48 ; 1.75] | -3.34<br>[-4.07 ; -2.63] | 0.0032*<br>[0.00025 ; 0.012] |
|  | PC2 | 9.48<br>[8.38 ; 10.55] | 1.62<br>[1.48 ; 1.76] | 5.88<br>[4.99 ; 6.81] | < 0.0001*<br>[< 0.0001 ; < 0.0001] |
|  | cos(moon) | -2.11<br>[-4.11 ; -0.22] | 1.63<br>[1.49 ; 1.79] | -1.31<br>[-2.61 ; -0.14] | 0.27<br>[0.013 ; 0.83] |

**Table S3.**
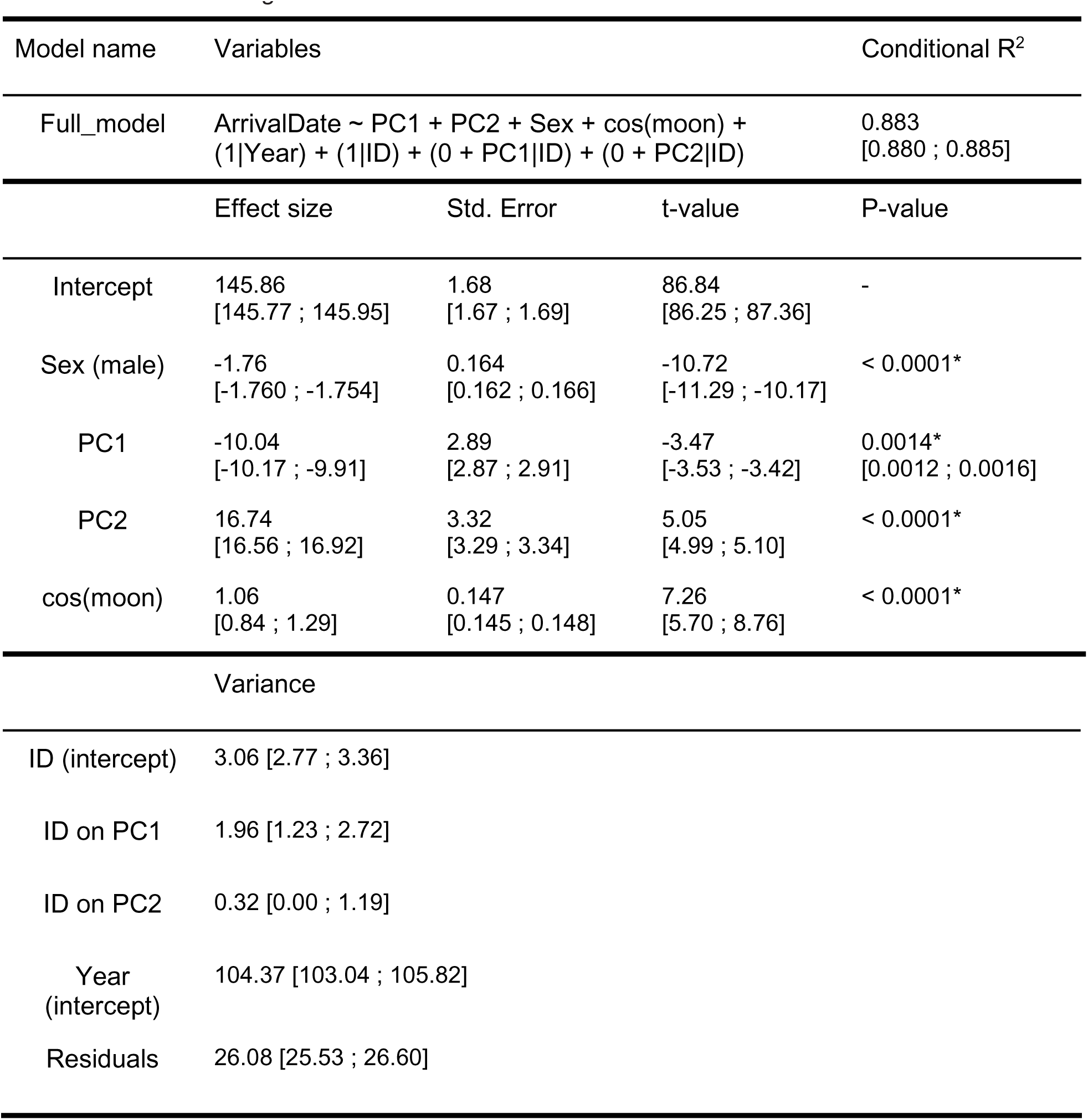
Detailed description of the model used to check for the effect of environmental variables on the phenology at the individual level. Sex is included to observe differences between males and females. The response variable ArrivalDate is a vector of dates of arrival at the breeding site for each individual over the study period. PC1 and PC2 are the first two components of the PCA performed on the climatic data. Cos(moon) is the cosine of the lunar angle for the arrival date. ID refers to the identity of each individual, and it is used as a random effect on both intercept and the slopes of PC1 and PC2. Finally, Year is included as a random effect to account for additional unexplained variation that might be caused by sampling variation. The second and third part of the table provide details on the estimates for the fixed and random effects respectively. The values shown are the mean value out of the 1000 models ran on simulated datasets and in square brackets we show the 2.5th and the 97.5th percentiles of each parameter. No p-values are provided for the intercept because this test is not of biological interest.

**Table S4.** Summary of the piecewise regressions on the five focal environmental variables. MinT_Spring is the average minimum daily spring temperature, MinT_Winter is the average minimum daily winter temperature, Prec_Spring is the total precipitation in spring, SWE_Spring is the total spring snow water equivalent and SWE_Winter is the total winter snow water equivalent. We rescaled year to obtain more intuitive intercept estimates (year 1980 = 0). Slope1 refers to the segment before the breakpoint and Slope2 refers to the segment after the breakpoint. We also show the statistics associated with the identified breakpoint. Asterisks next to the p-values show significance at the 0.05 level. The p-value for Slope2 is NA since standard asymptotics do not apply (Muggeo, 2008). No p-values are provided for the intercept because this test is not of biological interest.

|  |  | Estimates | Std. Error | t-value | p-value |
| --- | --- | --- | --- | --- | --- |
| MinT_Spring | Intercept | -0.82 | 0.30 | -2.76 | - |
|  | Slope1 | 0.07 | 0.02 | 3.33 | 0.002* |
|  | Slope2 | -0.05 | 0.05 | -0.87 | NA |
|  | Breakpoint | 2007 | 4.60 | -1.92 | 0.06 |
| MinT_Winter | Intercept | -8.88 | 0.80 | -11.10 | - |
|  | Slope1 | 0.24 | 0.14 | 1.76 | 0.09 |
|  | Slope2 | -0.03 | 0.03 | -1.07 | NA |
|  | Breakpoint | 1990 | 3.56 | -1.05 | 0.30 |
| Prec_Spring | Intercept | 41.35 | 6.90 | 6.00 | - |
|  | Slope1 | 2.26 | 1.91 | 1.18 | 0.24 |
|  | Slope2 | -0.30 | 0.18 | -1.68 | NA |
|  | Breakpoint | 1986 | 3.91 | -1.51 | 0.14 |
| SWE_Spring | Intercept | 3654.0 | 707.3 | 5.17 | - |
|  | Slope1 | -219.20 | 132.49 | -1.65 | 0.11 |
|  | Slope2 | 37.68 | 24.17 | 1.56 | NA |
|  | Breakpoint | 1990 | 3.55 | 2.23 | 0.03* |
| SWE_Winter | Intercept | 2643.3 | 511.4 | 5.17 | - |
|  | Slope1 | -156.42 | 107.41 | -1.46 | 0.15 |
|  | Slope2 | 28.31 | 15.93 | 1.78 | NA |
|  | Breakpoint | 1989 | 3.62 | 1.88 | 0.07 |

**Table S5.** Details of the five principal components. For each principal component we report its standard deviation, the proportion of variance explained and the cumulative proportion of this variance. In our linear regression we kept the first two principal components as their standard deviation is >1 (i.e. their eigenvalue >1) and combined they explain >70 % of the variance.

|  | PC1 | PC2 | PC3 | PC4 | PC5 |
| --- | --- | --- | --- | --- | --- |
| Standard deviation | 1.47 | 1.18 | 0.83 | 0.82 | 0.33 |
| Proportion of variance | 0.43 | 0.28 | 0.14 | 0.13 | 0.02 |
| Cumulative proportion | 0.43 | 0.71 | 0.85 | 0.98 | 1.00 |

**Table S6.** Loadings of the five original environmental variables from which the five principal components are constructed. MinT_Spring is the average minimum daily spring temperature, MinT_Winter is the average minimum daily winter temperature, Prec_Spring is the total precipitation in spring, SWE_Spring is the total spring snow water equivalent and SWE_Winter is the total winter snow water equivalent. PC1 is mainly driven by MinT_Winter, SWE_Spring and SWE_Winter, while PC2 mostly by MinT_Spring and Prec_Spring.

|  | PC1 | PC2 | PC3 | PC4 | PC5 |
| --- | --- | --- | --- | --- | --- |
| MinT_Spring | 0.08 | -0.68 | 0.33 | -0.65 | 0.09 |
| MinT_Winter | 0.45 | -0.19 | 0.64 | 0.58 | 0.07 |
| Prec_Spring | -0.04 | 0.68 | 0.60 | -0.41 | -0.06 |
| SWE_Spring | -0.64 | -0.05 | 0.20 | 0.17 | 0.72 |
| SWE_Winter | -0.61 | -0.22 | 0.27 | 0.19 | -0.69 |

## Notes

https://doi.org/10.5281/zenodo.7333319

### Competing Interest Statement

The authors have declared no competing interest.

### Summary of Updates

Added the Peer Community in Ecology Badge in the first page, as this paper has been recommended by them. Added a line in the acknowledgements to thank the reviewers and the recommender.

## References

Agostinelli, C., & Lund, U. (2017). R package ‘circular’: Circular statistics. https://r-forge.r-project.org/projects/circular/

Arietta, A. Z. A., Freidenburg, L. K., Urban, M. C., Rodrigues, S. B., Rubinstein, A., & Skelly, D. K. (2020). Phenological delay despite warming in wood frog *Rana sylvatica* reproductive timing: A 20-year study. Ecography, 43(12), 1791–1800. https://doi.org/10.1111/ecog.05297

Arnfield, H., Grant, R. A., Monk, C., & Uller, T. (2012). Factors influencing the timing of spring migration in common toads (*Bufo bufo*): Timing of spring migration in toads. Journal of Zoology, 288(2), 112–118. https://doi.org/10.1111/j.1469-7998.2012.00933.x

Barton, K. (2019). *Mult*i-Model Inference.

Beebee, T. J. C. (1995). Amphibian breeding and climate. Nature, 374(6519), 219–220. https://doi.org/10.1038/374219a0

Begert, M., & Frei, C. (2018). Long-term area-mean temperature series for Switzerland—Combining homogenized station data and high resolution grid data. International Journal of Climatology, 38(6), 2792–2807. https://doi.org/10.1002/joc.5460

Bison, M., Yoccoz, N. G., Carlson, B. Z., Klein, G., Laigle, I., Van Reeth, C., Asse, D., & Delestrade, A. (2020). Best environmental predictors of breeding phenology differ with elevation in a common woodland bird species. Ecology and Evolution, 10(18), 10219–10229. https://doi.org/10.1002/ece3.6684

Bison, M., Yoccoz, N. G., Carlson, B. Z., Klein, G., Laigle, I., Van Reeth, C., & Delestrade, A. (2021). Earlier snowmelt advances breeding phenology of the common frog (*Rana temporaria*) but increases the risk of frost exposure and wetland drying. Frontiers in Ecology and Evolution, 9, 645585. https://doi.org/10.3389/fevo.2021.645585

Blaustein, A. R., Belden, L. K., Olson, D. H., Green, D. M., Root, T. L., & Kiesecker, J. M. (2001). Amphibian breeding and climate change. Conservation Biology, 15(6), 1804–1809. https://doi.org/10.1046/j.1523-1739.2001.00307.x

Bundesamt für Umwelt (BAFU). (2020). Klimawandel in der Schweiz (p. 105).

Clermont, J., Réale, D., & Giroux, J.-F. (2018). Plasticity in laying dates of Canada geese in response to spring phenology. Ibis, 160(3), 597–607. https://doi.org/10.1111/ibi.12560

Corn, P. S. (2003). Amphibian breeding and climate change: Importance of snow in the mountains. Conservation Biology, 17(2), 622–625. https://doi.org/10.1046/j.1523-1739.2003.02111.x

Corn, P. S., & Muths, E. (2002). Variable breeding phenology affects the exposure of amphibian embryos to ultraviolet radiation. Ecology, 83(11), 2958–2963. https://doi.org/10.1890/0012-9658(2002)083[2958:VBPATE]2.0.CO;2

Diaz, H. F., Grosjean, M., & Graumlich, L. (2003). Climate variability and change in high elevation regions: Past, present and future. Climatic Change, 59, 1–4. https://doi.org/10.1007/978-94-015-1252-7_1

Duellman, W. E., & Trueb, L. (1986). Biology of Amphibians. McGraw-Hill Book Company.

Falconer, D. S. (1981). Introduction to quantitative genetics (2d ed.). Longman.

Feldmeier, S., Schmidt, B. R., Zimmermann, N. E., Veith, M., Ficetola, G. F., & Lötters, S. (2020). Shifting aspect or elevation? The climate change response of ectotherms in a complex mountain topography. Diversity and Distributions, 26(11), 1483–1495. https://doi.org/10.1111/ddi.13146

Ficetola, G. F., & Maiorano, L. (2016). Contrasting effects of temperature and precipitation change on amphibian phenology, abundance and performance. Oecologia, 181(3), 683–693. https://doi.org/10.1007/s00442-016-3610-9

Fitak, R. R., & Johnsen, S. (2017). Bringing the analysis of animal orientation data full circle: Model-based approaches with maximum likelihood. Journal of Experimental Biology, 220(21), 3878–3882. https://doi.org/10.1242/jeb.167056

Franklin, K. A., Nicoll, M. A. C., Butler, S. J., Norris, K., Ratcliffe, N., Nakagawa, S., & Gill, J. A. (2022). Individual repeatability of avian migration phenology: A systematic review and meta-analysis. Journal of Animal Ecology, 91(7), 1416–1430. https://doi.org/10.1111/1365-2656.13697

Garner, T. W. J., Rowcliffe, J. M., & Fisher, M. C. (2011). Climate change, chytridiomycosis or condition: An experimental test of amphibian survival. Global Change Biology, 17(2), 667–675. https://doi.org/10.1111/j.1365-2486.2010.02272.x

Gelman, A. (2008). Scaling regression inputs by dividing by two standard deviations. Statistics in Medicine, 27(15), 2865–2873. https://doi.org/10.1002/sim.3107

Gittins, S. P., Parker, A. G., & Slater, F. M. (1980). Population characteristics of the common toad (*Bufo bufo*) visiting a breeding site in Mid-Wales. Journal of Animal Ecology, 49(1), 161–173. https://doi.org/10.2307/4281

Gotthard, K. (2001). Growth strategies of ectothermic animals in temperate environments. In D. Atkinson & M. Thorndyke (Eds.), Environment and animal development: Genes, life histories and plasticity (pp. 287–304). BIOS Scientific.

Grant, R. A., Chadwick, E. A., & Halliday, T. R. (2009). The lunar cycle: A cue for amphibian reproductive phenology? Animal Behaviour, 78(2), 349–357. https://doi.org/10.1016/j.anbehav.2009.05.007

Green, D. M. (2017). Amphibian breeding phenology trends under climate change: Predicting the past to forecast the future. Global Change Biology, 23(2), 646–656. https://doi.org/10.1111/gcb.13390

Green, T., Das, E., & Green, D. M. (2016). Springtime emergence of overwintering toads, *Anaxyrus fowleri*, in relation to environmental factors. Copeia, 104(2), 393–401. https://doi.org/10.1643/CE-15-323

Grossenbacher, K. (2002). First results of a 20-year-study on common toad *Bufo bufo* in the Swiss Alps. Biota, 3(1–2), 43–48.

Hartel, T., Sas-Kovacs, I., Pernetta, A., & Geltsch, I. (2007). The reproductive dynamics of temperate amphibians: A review. North-Western Journal of Zoology, 3.

Hemelaar, A. (1988). Age, growth and other population characteristics of *Bufo bufo* from different latitudes and altitudes. Journal of Herpetology, 22(4), 369. https://doi.org/10.2307/1564332

Heusser, H., & Ott, J. (1968). Sollzeit der Laichwanderung bei der Erdkröte, Bufo bufo (L.). Revue Suisse de Zoologie, 75, 1005–1022.

Höglund, J., & Robertson, J. G. M. (1987). Random mating by size in a population of common toads (*Bufo bufo*). Amphibia-Reptilia, 8(4), 321–330. https://doi.org/10.1163/156853887X00108

Höglund, J., & Robertson, J. G. M. (1988). Chorusing behaviour, a density-dependent alternative mating strategy in male common toads (*Bufo bufo*). Ethology, 79(4), 324–332. https://doi.org/10.1111/j.1439-0310.1988.tb00721.x

Iler, A. M., CaraDonna, P. J., Forrest, J. R. K., & Post, E. (2021). Demographic consequences of phenological shifts in response to climate change. Annual Review of Ecology, Evolution, and Systematics, 52(1), annurev-ecolsys-011921-032939. https://doi.org/10.1146/annurev-ecolsys-011921-032939

Ims, R. A. (1990). The ecology and evolution of reproductive synchrony. Trends in Ecology & Evolution, 5(5), 135–140. https://doi.org/10.1016/0169-5347(90)90218-3

Jara, F. G., Thurman, L. L., Montiglio, P.-O., Sih, A., & Garcia, T. S. (2019). Warming-induced shifts in amphibian phenology and behavior lead to altered predator–prey dynamics. Oecologia, 189(3), 803–813. https://doi.org/10.1007/s00442-019-04360-w

Jarvis, L. E., Grant, R. A., & SenGupta, A. (2021). Lunar phase as a cue for migrations to two species of explosive breeding amphibians—Implications for conservation. European Journal of Wildlife Research, 67(1), 11. https://doi.org/10.1007/s10344-020-01453-3

Keiler, M., Knight, J., & Harrison, S. (2010). Climate change and geomorphological hazards in the eastern European Alps. *Philosophical Transactions of the Royal Society A: Mathematical*, Physical and Engineering Sciences, 368(1919), 2461–2479. https://doi.org/10.1098/rsta.2010.0047

Kokko, H. (1999). Competition for early arrival in migratory birds. Journal of Animal Ecology, 68(5), 940–950. https://doi.org/10.1046/j.1365-2656.1999.00343.x

Körner, C., & Hiltbrunner, E. (2021). Why is the alpine flora comparatively robust against climatic warming? Diversity, 13(8), 383. https://doi.org/10.3390/d13080383

Kovar, R., Brabec, M., Bocek, R., & Vita, R. (2009). Spring migration distances of some Central European amphibian species. Amphibia-Reptilia, 30(3), 367–378. https://doi.org/10.1163/156853809788795236

Kürten, N., Schmaljohann, H., Bichet, C., Haest, B., Vedder, O., González-Solís, J., & Bouwhuis, S. (2022). High individual repeatability of the migratory behaviour of a long-distance migratory seabird. Movement Ecology, 10(1), 5. https://doi.org/10.1186/s40462-022-00303-y

Kuznetsova, A., Brockhoff, P. B., & Christensen, R. H. B. (2017). lmerTest Package: Tests in linear mixed effects models. Journal of Statistical Software, 82(13), 1–26.

Landler, L., Ruxton, G. D., & Malkemper, E. P. (2018). Circular data in biology: Advice for effectively implementing statistical procedures. Behavioral Ecology and Sociobiology, 72(8), 128. https://doi.org/10.1007/s00265-018-2538-y

Lazaridis, E. (2014). *lunar: Lunar phase and distance, seasons and other environmental factors* (Version 0.1-04).

Lessells, C. M., & Boag, P. T. (1987). Unrepeatable repeatabilities: A common mistake. The Auk, 104(1), 116–121. https://doi.org/10.2307/4087240

Loman, J., & Madsen, T. (1986). Reproductive tactics of large and small male toads *Bufo bufo*. Oikos, 46(1), 57. https://doi.org/10.2307/3565380

Lüscher, B., Beer, S., & Grossenbacher, K. (2016). Die Höhenverbreitung der Erdkröte (*Bufo bufo*) im Berner Oberland (Schweiz) unter sich verändernden Klimabedingungen. Zeitschrift für Feldherpetologie, 23, 47–58.

Miller, D. A. W., Grant, E. H. C., Muths, E., Amburgey, S. M., Adams, M. J., Joseph, M. B., Waddle, J. H., Johnson, P. T. J., Ryan, M. E., Schmidt, B. R., Calhoun, D. L., Davis, C. L., Fisher, R. N., Green, D. M., Hossack, B. R., Rittenhouse, T. A. G., Walls, S. C., Bailey, L. L., Cruickshank, S. S., … Sigafus, B. H. (2018). Quantifying climate sensitivity and climate-driven change in North American amphibian communities. Nature Communications, 9(1), 3926. https://doi.org/10.1038/s41467-018-06157-6

Morin, P. J., Lawler, S. P., & Johnson, E. A. (1990). Ecology and breeding phenology of larval *Hyla andersonii*: The disadvantages of breeding late. Ecology, 71(4), 1590–1598. https://doi.org/10.2307/1938294

Muggeo, V. M. R. (2008). Segmented: An R package to fit regression models with broken-line. R News, 8(1), 20–25.

Muir, A. P., Biek, R., Thomas, R., & Mable, B. K. (2014). Local adaptation with high gene flow: Temperature parameters drive adaptation to altitude in the common frog (*Rana temporaria*). Molecular Ecology, 23(3), 561–574. https://doi.org/10.1111/mec.12624

National Academies of Sciences, Engineering, and Medicine. (2016). Attribution of extreme weather events in the context of climate change. National Academies Press.

Nogués-Bravo, D., Araújo, M. B., Errea, M. P., & Martínez-Rica, J. P. (2007). Exposure of global mountain systems to climate warming during the 21st Century. Global Environmental Change, 17(3–4), 420–428. https://doi.org/10.1016/j.gloenvcha.2006.11.007

Nufio, C. R., McGuire, C. R., Bowers, M. D., & Guralnick, R. P. (2010). Grasshopper community response to climatic change: Variation along an elevational gradient. PLoS ONE, 5(9), 1–11. https://doi.org/10.1371/journal.pone.0012977

Orizaola, G., Richter-Boix, A., & Laurila, A. (2016). Transgenerational effects and impact of compensatory responses to changes in breeding phenology on antipredator defenses. Ecology, 97(9), 2470–2478. https://doi.org/10.1002/ecy.1464

Oseen, K. L., & Wassersug, R. J. (2002). Environmental factors influencing calling in sympatric anurans. Oecologia, 133(4), 616–625. https://doi.org/10.1007/s00442-002-1067-5

Parmesan, C. (2007). Influences of species, latitudes and methodologies on estimates of phenological response to global warming. Global Change Biology, 13(9), 1860–1872. https://doi.org/10.1111/j.1365-2486.2007.01404.x

Parmesan, C., & Yohe, G. (2003). A globally coherent fingerprint of climate change impacts across natural systems. Nature, 421(6918), 37–42. https://doi.org/10.1038/nature01286

Phillimore, A. B., Hadfield, J. D., Jones, O. R., & Smithers, R. J. (2010). Differences in spawning date between populations of common frog reveal local adaptation. Proceedings of the National Academy of Sciences, 107(18), 8292–8297. https://doi.org/10.1073/pnas.0913792107

Pollock, K. H. (1982). A capture-recapture design robust to unequal probability of capture. The Journal of Wildlife Management, 46(3), 752–757. https://doi.org/10.2307/3808568

Prodon, R., Geniez, P., Cheylan, M., & Besnard, A. (2020). Amphibian and reptile phenology: The end of the warming hiatus and the influence of the NAO in the North Mediterranean. International Journal of Biometeorology, 64(3), 423–432. https://doi.org/10.1007/s00484-019-01827-6

Quinn, J. A., & Wetherington, J. D. (2002). Genetic variability and phenotypic plasticity in flowering phenology in populations of two grasses. The Journal of the Torrey Botanical Society, 129(2), 96–106. https://doi.org/10.2307/3088723

R Core Team. (2020). R: A language and environment for statistical computing. R foundation for statistical computing, Vienna, Austria. URL https://www.R-project.org/.

R Studio Team. (2022). RStudio: Integrated Development for R. RStudio, PBC., Boston, MA

Rahmstorf, S., & Coumou, D. (2011). Increase of extreme events in a warming world. Proceedings of the National Academy of Sciences, 108(44), 17905–17909. https://doi.org/10.1073/pnas.1101766108

Reading, C. J. (2003). The effects of variation in climatic temperature (1980–2001) on breeding activity and tadpole stage duration in the common toad, *Bufo bufo*. Science of The Total Environment, 310(1–3), 231–236. https://doi.org/10.1016/S0048-9697(02)00643-5

Reading, C. J. (2007). Linking global warming to amphibian declines through its effects on female body condition and survivorship. Oecologia, 151(1), 125–131. https://doi.org/10.1007/s00442-006-0558-1

Reading, C. J. (2010). The impact of environmental temperature on larval development and metamorph body condition in the common toad, *Bufo bufo*. Amphibia-Reptilia, 31(4), 483–488. https://doi.org/10.1163/017353710X521537

Reading, C. J., & Clarke, R. T. (1983). Male breeding behaviour and mate acquisition in the common toad, *Bufo bufo*. Journal of Zoology, 201(2), 237–246. https://doi.org/10.1111/j.1469-7998.1983.tb04273.x

Reading, C. J., & Clarke, R. T. (1999). Impacts of climate and density on the duration of the tadpole stage of the common toad *Bufo bufo*. Oecologia, 121(3), 310–315. https://doi.org/10.1007/s004420050933

Rebetez, M., & Reinhard, M. (2008). Monthly air temperature trends in Switzerland 1901-2000 and 1975–2004. Theoretical and Applied Climatology, 91(1), 27–34. https://doi.org/10.1007/s00704-007-0296-2

Reinhardt, T., Steinfartz, S., & Weitere, M. (2015). Inter-annual weather variability can drive the outcome of predator prey match in ponds. Amphibia-Reptilia, 36(2), 97–109. https://doi.org/10.1163/15685381-00002982

Scherrer, D., & Körner, C. (2011). Topographically controlled thermal-habitat differentiation buffers alpine plant diversity against climate warming. Journal of Biogeography, 38(2), 406–416. https://doi.org/10.1111/j.1365-2699.2010.02407.x

Semlitsch, R. D. (1985). Analysis of climatic factors influencing migrations of the salamander *Ambystoma talpoideum*. Copeia, 1985(2), 477–489. https://doi.org/10.2307/1444862

Semlitsch, R. D., Scott, D. E., Pechmann, J. H. K., & Gibbons, J. W. (1993). Phenotypic variation in the arrival time of breeding salamanders: Individual repeatability and environmental influences. The Journal of Animal Ecology, 62(2), 334. https://doi.org/10.2307/5364

Sinsch, U. (1988). Seasonal changes in the migratory behaviour of the toad *Bufo bufo*: Direction and magnitude of movements. Oecologia, 76, 390–398.

Sinsch, U., & Schäfer, A. M. (2016). Density regulation in toad populations (*Epidalea calamita*, Bufotes viridis) by differential winter survival of juveniles. Journal of Thermal Biology, 55, 20–29. https://doi.org/10.1016/j.jtherbio.2015.11.007

Stoffel, M. A., Nakagawa, S., & Schielzeth, H. (2017). RptR: repeatability estimation and variance decomposition by generalized linear mixed-effects models. Methods in Ecology and Evolution, 8, 1639–1644. https://doi.org/doi:10.1111/2041-210X.12797

Sztatecsny, M., & Schabetsberger, R. (2005). Into thin air: Vertical migration, body condition, and quality of terrestrial habitats of alpine common toads, *Bufo bufo*. Canadian Journal of Zoology, 83, 788–796.

Tang, J., Körner, C., Muraoka, H., Piao, S., Shen, M., Thackeray, S. J., & Yang, X. (2016). Emerging opportunities and challenges in phenology: A review. Ecosphere, 7(8), e01436. https://doi.org/10.1002/ecs2.1436

Thompson, L. (2000). Ice core evidence for climate change in the Tropics: Implications for our future. Quaternary Science Reviews, 19, 19–35. https://doi.org/10.1016/S0277-3791(99)00052-9

Thornton, P. E., Running, S. W., & White, M. A. (1997). Generating surfaces of daily meteorological variables over large regions of complex terrain. Journal of Hydrology, 190(3), 214–251. https://doi.org/10.1016/S0022-1694(96)03128-9

Todd, B., Scott, D., Pechmann, J., & Gibbons, J. (2011). Climate change correlates with rapid delays and advancements in reproductive timing in an amphibian community. Proceedings. Biological Sciences / The Royal Society, 278, 2191–2197. https://doi.org/10.1098/rspb.2010.1768

Tryjanowski, P., Rybacki, M., & Sparks, T. (2003). Changes in the first spawning dates of common frogs and common toads in western Poland in 1978–2002. Annales Zoologici Fennici, 40, 459–464.

Turner, R. K., & Maclean, I. M. D. (2022). Microclimate-driven trends in spring-emergence phenology in a temperate reptile (*Vipera berus*): Evidence for a potential “climate trap”? Ecology and Evolution, 12(2), e8623. https://doi.org/10.1002/ece3.8623

Urban, M. C. (2018). Escalator to extinction. Proceedings of the National Academy of Sciences, 115(47), 11871–11873. https://doi.org/10.1073/pnas.1817416115

Vaillant, J. L., Potti, J., Camacho, C., Canal, D., & Martínez-Padilla, J. (2021). Low repeatability of breeding events reflects flexibility in reproductive timing in the pied flycatcher *Ficedula hypoleuca* in Spain. Ardeola, 69(1), 21–39. https://doi.org/10.13157/arla.69.1.2022.ra2

Visser, M. E., & Gienapp, P. (2019). Evolutionary and demographic consequences of phenological mismatches. Nature Ecology & Evolution, 3(6), 879–885. https://doi.org/10.1038/s41559-019-0880-8

Vitasse, Y., Ursenbacher, S., Klein, G., Bohnenstengel, T., Chittaro, Y., Delestrade, A., Monnerat, C., Rebetez, M., Rixen, C., Strebel, N., Schmidt, B. R., Wipf, S., Wohlgemuth, T., Yoccoz, N. G., & Lenoir, J. (2021). Phenological and elevational shifts of plants, animals and fungi under climate change in the European Alps. Biological Reviews, 96(5), 1816–1835. https://doi.org/10.1111/brv.12727

Wells, K. D. (1977). The social behaviour of anuran amphibians. Animal Behaviour, 25, 666–693. https://doi.org/10.1016/0003-3472(77)90118-X

While, G. M., & Uller, T. (2014). Quo vadis amphibia? Global warming and breeding phenology in frogs, toads and salamanders. Ecography, 37(10), 921–929. https://doi.org/10.1111/ecog.00521

Zani, P. A. (2008). Climate change trade-offs in the side-blotched lizard (Uta stansburiana): Effects of growing-season length and mild temperatures onwinter survival. Physiological and Biochemical Zoology, 81(6), 797–809. https://doi.org/10.1086/588305

Zeileis, A., & Grothendieck, G. (2005). zoo: S3 infrastructure for regular and irregular time series. Journal of Statistical Software, 14, 1–27. https://doi.org/10.18637/jss.v014.i06

